# Reproducible switching between a walled and cell wall-deficient lifestyle of actinomycetes using gradient agar plates

**DOI:** 10.1101/2022.11.14.516409

**Authors:** Maarten Lubbers, Gilles P. van Wezel, Dennis Claessen

## Abstract

The cell wall is a shape-defining structure that envelopes almost all bacteria, protecting them from biotic and abiotic stresses. Paradoxically, some filamentous actinomycetes have a natural ability to shed their cell wall under influence of hyperosmotic stress. These wall-deficient cells can revert to their walled state when transferred to a medium without osmoprotection but often lyse due to their fragile nature. Here, we designed plates with an osmolyte gradient to reduce cell lysis and thereby facilitating the transition between a walled and wall-deficient state. These gradient plates allow determining of the osmolyte concentration where switching takes place, thereby enabling careful and reproducible comparison between mutants affected by switching. Exploring these transitions could give valuable insights into the ecology of actinomycetes and their biotechnological applications.

**HIGHLIGHTS:**

- Using agar plates with a gradient of osmoprotectants, revertant Streptomycetes can gradually revert to a walled state, thereby dramatically decreasing lysis.
- Our method allows precise determination of the osmolyte concentration where reversion takes place, allowing careful and reproducible comparison between mutants.
- Gradient agar plates can also be used to study chemical differentiation in Streptomycetes as a response to osmotic stress.

## 1. INTRODUCTION

Bacteria occur in a plethora of ecosystems, ranging from high mountains to deep-sea vents. In these habitats, they are challenged by environmental fluctuations, such as changes in pH, temperature, and water availability (Nguyen et al., 2021). These fluctuations are important cues for morphological and chemical differentiation. For instance, bacteria can adapt to osmotic stress conditions, such as desiccation and hypertonicity, by modulating their fatty acid composition in the membrane, synthesizing and accumulating osmoprotectants, or secreting extracellular polymeric substances, amongst others (Maccario et al., 2015; de Maayer et al., 2014; Lebre et al., 2017). Recently, another response to hyperosmotic stress was shown that leads to a dramatic change in morphology. More specifically, some filamentous Actinobacteria were found to extrude wall-less deficient cells as an adaptation to hyperosmotic stress conditions (Ramijan et al., 2018). These so-called S-cells (for Stress-induced cells) are extruded from hyphal tips into the environment and require high levels of osmolytes to prevent them from bursting (Ramijan et al., 2018). Interestingly, recent evidence suggests that such wall-deficient cells provide protection against bacteriophage attack, indicating that they could play a central role in the evolutionary arms race between phages and bacteria (Ongenae et al., 2022). Over time, S-cells increase in size but do not proliferate without their cell wall. However, prolonged exposure of S-cells to hyperosmotic stress can lead to the formation of so-called L-forms, which are wall-deficient cells that do propagate without their cell wall (Ramijan et al., 2018). Both S-cells and L-forms can switch to the walled state, which is stimulated by removing the protective osmolytes from the environment. This so-called reversible metamorphosis provides unprecedented and innovative leads to study the requirements for filamentous and wall-less growth.

One challenge when working with L-forms has been to find the conditions that allow predictable and reproducible switching from the wall-deficient to the walled state. For instance, *Kitasatospora viridifaciens* L-forms are currently triggered to switch to the filamentous mode of growth by plating high densities of L-form cells on solid media without osmoprotectants (Zhang et al., 2021). In this case, only a small fraction of cells will manage to switch to the walled state. However, for many mutant strains, it has been found challenging or impossible to revert to a walled state: as wall-deficient cells are fragile, there is a high chance of bursting after such a dramatic change in medium composition. Furthermore, current methods cannot rapidly determine the osmolyte concentration at which switching takes place, which would provide a careful and reproducible comparison between mutants. Gradient agar plates are a powerful technique originally developed in the 1950s, allowing researchers to measure antibiotic resistance or response to a substance of interest (Szybalski and Bryson, 1952). We here describe the application of such gradient agar plates to study growth on media with an osmolyte gradient, which provides rapid insights into switching efficiencies at various osmolyte concentrations. Using *K. viridifaciens* L-forms as an example, we show that this allows us to improve the switching frequency from the wall-deficient to walled state more than 2-fold, while also providing rapid insights into the switching efficiency of different mutants. Altogether, these results demonstrate the power of such plates for studying reversible metamorphosis in bacteria.

## 2. MATERIAL AND METHODS

### 2.1 Strains and media

The strains used in this study are shown in Table 1. The reverting *K. viridifaciens* L-form strains *alpha* (Ramijan et al., 2018) and *delta* (Shitut et al., 2022) were obtained after induction with penicillin and lysozyme. Strain *K. viridifaciens* M1 is a natural L-form cell line, obtained after prolonged exposure of S-cells to hyperosmotic stress (Ramijan et al., 2018). The non-switching L-form strain *K. viridifaciens* Δ*dcw* was created by replacing *ftsW, murG, ftsQ, ftsZ, ylmD, ylmE, sepG, sepF*, and *divIVA* with the apramycin resistance marker *aac(3)IV* in *alpha* (Zhang et al., 2021). To obtain spores, *K. viridifaciens* DSM40239 and *Streptomyces coelicolor* A3(2) M145 were grown at 30°C for 4 days on Maltose-Yeast Extract-Malt Extract (MYM) medium, after which spores were harvested as described (Kieser et al., 2000). The spores were then used as an inoculum for TSBS cultures (tryptone soy broth with 10% sucrose (Kieser et al., 2000)), which were grown for 3 days in flasks equipped with a metal coil. L-forms strains were grown for 4 days in liquid L-phase broth (LPB) supplemented with 25 mM MgCl_2_ (Ramijan et al., 2018).

**Table 1.** Strains used in this study.

| Strains | Genotype | Reference |
| --- | --- | --- |
| *Kitasatospora viridifaciens* DSM40239 | Wild-type | DSMZ (Ramijan et al., 2017) |
| *Streptomyces coelicolor* A3(2) M145 | Wild-type | Lab collection |
| *S. coelicolor* $\Delta bldB$ | Mutant | Lab collection (Pope et al., 1998) |
| *S. coelicolor* $\Delta bldC$ | Mutant | Lab collection (Hunt et al., 2005) |
| *S. coelicolor* $\Delta bldF$ | Mutant | Lab collection (Puglia et al., 1984) |
| *S. coelicolor* $\Delta actVB$ | Mutant | Lab collection |
| *S. coelicolor* $\Delta actII-ORF4$ | Mutant | Lab collection |
| *K. viridifaciens* *alpha* | Mutant | Lab collection (Ramijan et al., 2018) |
| *K. viridifaciens* *delta* | Mutant | Lab collection (Shitut et al., 2021) |
| *K. viridifaciens* M1 | Mutant | Lab collection (Ramijan et al., 2018) |
| *K. viridifaciens* $\Delta dcw$ | Mutant | Lab collection (Ramijan et al., 2019) |

### 2.2 Gradient agar plates

To create gradient agar plates, 50 mL of non-osmoprotective MYM medium (Stuttard, 1982) was poured into a 120×120 mm square petri dish (Greiner Bio-One), after which the plates were allowed to settle under a 5° angle. After the medium was solidified, 40 mL of osmoprotective LPMA medium supplemented with 5% (v/v) horse serum and 25 mM MgCl_2_ (Ramijan et al., 2018) was poured on top of the MYM medium (Sup. Fig. 1). Importantly, the LPMA was poured from the thinnest side of the MYM medium to ensure that the gradient ends in a straight line. This is in contrast to most other described gradient plate methods that typically rely on an equal volume of both media types (Szybalski and Bryson, 1952; Sacks, 1956). In this study, however, the two opposite ends of the plate should consist of only one medium type as this allows for a clear comparison between LPMA, MYM, and the gradient.

**Figure 1.**
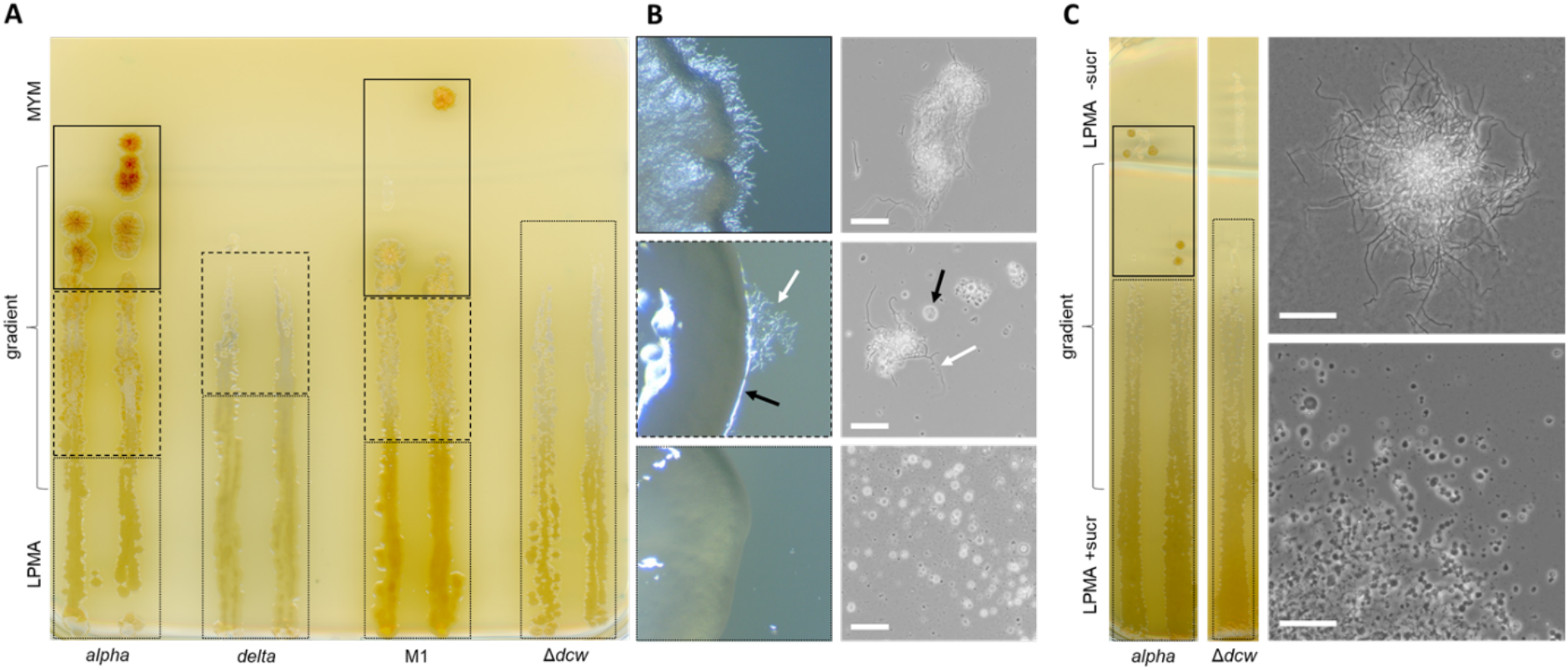
*K. viridifaciens* L-form strains display morphological differentiation on gradient agar plates. (**A**). Morphological transitions on MYM/LPMA gradient plates. Strains *alpha* and M1 show filamentous (solid line), heteromorphic (dashed line), or wall-less growth (dotted line) depending on the osmolyte concentration. By contrast, *delta* shows either wall-less (dotted line) or heteromorphic growth (dashed line). Δ*dcw* only proliferates as wall-less cells. (**B**) Heteromorphic colonies of *alpha* show both filamentous (white arrow) and wall-less (black arrow) growth on the edges of colonies. (**C**). In comparison to MYM/LPMA gradient plates, Δ*dcw* shows an identical transition pattern on plates containing a gradient of sucrose. However, unlike on LPMA/MYM gradient plates, the heteromorphic state is absent in *alpha* on LPMA gradient plates in which only the sucrose concentration is varied. The scale bar represents 40 µm.

To monitor growth on gradient agar plates, strains were pre-grown in liquid media. To this end, spores or mycelium of *S. coelicolor* A3(2) M145, *S. coelicolor* Δ*actII-ORF4, S. coelicolor* Δ*actVB, S. coelicolor* Δ*bldB, S. coelicolor* Δ*bldC, S. coelicolor* Δ*bldF*, or *K. viridifaciens* DSM40239 were inoculated and grown for 3 days in coiled flasks in TSBS medium (25 ml) at 30°C while shaking at 200 rpm, after which mycelium was transferred to the gradient agar plates using an inoculation loop. Likewise, L-form cultures were grown in L-phase Broth (LPB) medium (Ramijan et al., 2018) in a shaker at 30°C at 100 rpm for 3 days and subsequently transferred to gradient agar plates using an inoculation loop. Gradient agar plates were sealed with parafilm and incubated at 30 °C. As we assume that the gradient is linear (Fig. S1), the percentage can be calculated as follows: *%LPMA = L*_*a-b*_ */ L*_*a-c*_ *x 100* (where a-c is the total length of the gradient in cm, of which a is 100% LPMA, and b is the location in the gradient for which the percentage is to be calculated).

### 2.3 Quantification of reversion efficiency

To investigate at which concentration most efficient switching from the wall-deficient to the walled state occurs, *alpha* was grown in LPB medium for 4 days at 30 °C while shaking at 100 rpm in an orbital shaker. At an OD_600_ of 1.4, 200 µL of the culture was confluently plated on plates containing mixtures of LPMA and MYM (ranging from 100:0 to 0:100 [MYM:LPMA] in steps of 20 percent). After 2 days of growth at 30 °C, the number of walled colonies was counted per plate using a PBI colony counter (Quartz). Experiments were performed in triplicate. Data were analyzed using GraphPad Prism (version 9.4.1).

### 2.4 Imaging

Gradient agar plates and 24-well plates were scanned using an Epson Perfection V600 Photo scanner. Colonies were photographed using a Mikrocam SP 5.0 microscope camera (Bresser) connected to a SteREO Discovery V8 stereomicroscope. Cells were imaged using a Zeiss Axio Lab A1 upright Microscope, equipped with an Axiocam MRc (Zeiss).

## 3. RESULTS AND DISCUSSION

### 3.1 Filamentous Actinobacteria display morphological differentiation on gradient agar plates

We previously showed that *K. viridifaciens alpha*, an unstable L-form strain obtained after induction with penicillin and lysozyme, can either grow filamentous or wall-deficient depending on the osmolarity of the media (Ramijan et al., 2018; Zhang et al., 2021). Using gradient plates, we indeed found that *alpha* could readily switch between these distinct morphologies (Fig. 1A). On 0-40% LPMA, *alpha* solely grew filamentous. It is important to note that in contrast to the parental strain *K. viridifaciens* DSM40239, *alpha* is unable to differentiate and form aerial hyphae (Zhang et al., 2021) (see also Fig. 3). Conversely, in the range of 90-100% LPMA, no filaments were detected, and proliferation of *alpha* occurred exclusively via the formation of wall-deficient cells. Interestingly, in between these zones (i.e. in the range of 40-90% LPMA), heteromorphic colonies were formed that consisted of a mixture of filaments and wall-deficient cells. Mycelium thereby protruded from the edges of the colonies, as revealed by light microscopy (Fig. 1B).

To ascertain that these morphological transitions on gradient plates are not unique for *alpha*, we also tested other L-form cell lines. Strain M1, a natural L-form cell line acquired after prolonged exposure of S-cells to hyperosmotic stress (Ramijan et al., 2018), showed similar transitions as *alpha* at comparable places on the gradient plates. More specifically, walled cells were formed during growth on 0-35% LPMA, whereas heteromorphic colonies were found in the range of 35-85% LPMA. Colonies consisting exclusively of wall-deficient cells were observed when using 85-100% LPMA. Contrary to the two other L-form strains, *K. viridifaciens delta*, which was obtained after induction with penicillin and lysozyme (Shitut et al., 2022) failed to grow in a canonical fashion on MYM. Instead, at 30-70% LPMA, heteromorphic colonies were formed. As expected, *K. viridifaciens* Δ*dcw*, a mutant L-form strain lacking a major part of the so-called *dcw* gene cluster (Zhang et al., 2021), failed to revert to its walled state (Fig. 1A). Interestingly, Δ*dcw* formed colonies consisting of L-form cells at 25% LPMA, apparently tolerating a lower concentration of osmoprotectants as compared to *alpha* and M1, both of which formed mycelial colonies at this concentration.

To study solely the effect of sucrose, alpha and Δ*dcw* were grown on LPMA gradient plates in which only the sucrose concentration was varied (Fig. 1B). Δ*dcw* showed an almost identical transition pattern as observed on the MYM/LPMA gradient plates. More specifically, growth was observed in the range of 25-100% sucrose. However, the transition pattern of *alpha* was different. Unlike on MYM/LPMA gradient plates, the heteromorphic state was absent. Walled growth was observed at 0-30% sucrose, while all colonies were wall-deficient above 30%. As gradients of LPMA and MYM gave more valuable information on morphological transitions, these plates were used for further experiments.

### 3.2 Gradient agar plates allow prediction of switching efficiency

To investigate whether morphological changes observed on gradient plates correspond to those in well-mixed plates, strains *alpha* and Δ*dcw* were grown in 24-well plates containing different LPMA:MYM ratios for three days at 30 °C. As expected, Δ*dcw* cells failed to grow at concentrations below 25% LPMA, as indicated by the absence of cells in well D1 (Fig. 2A), which are similar to those observed on gradient agar plates.

**Figure 2.**
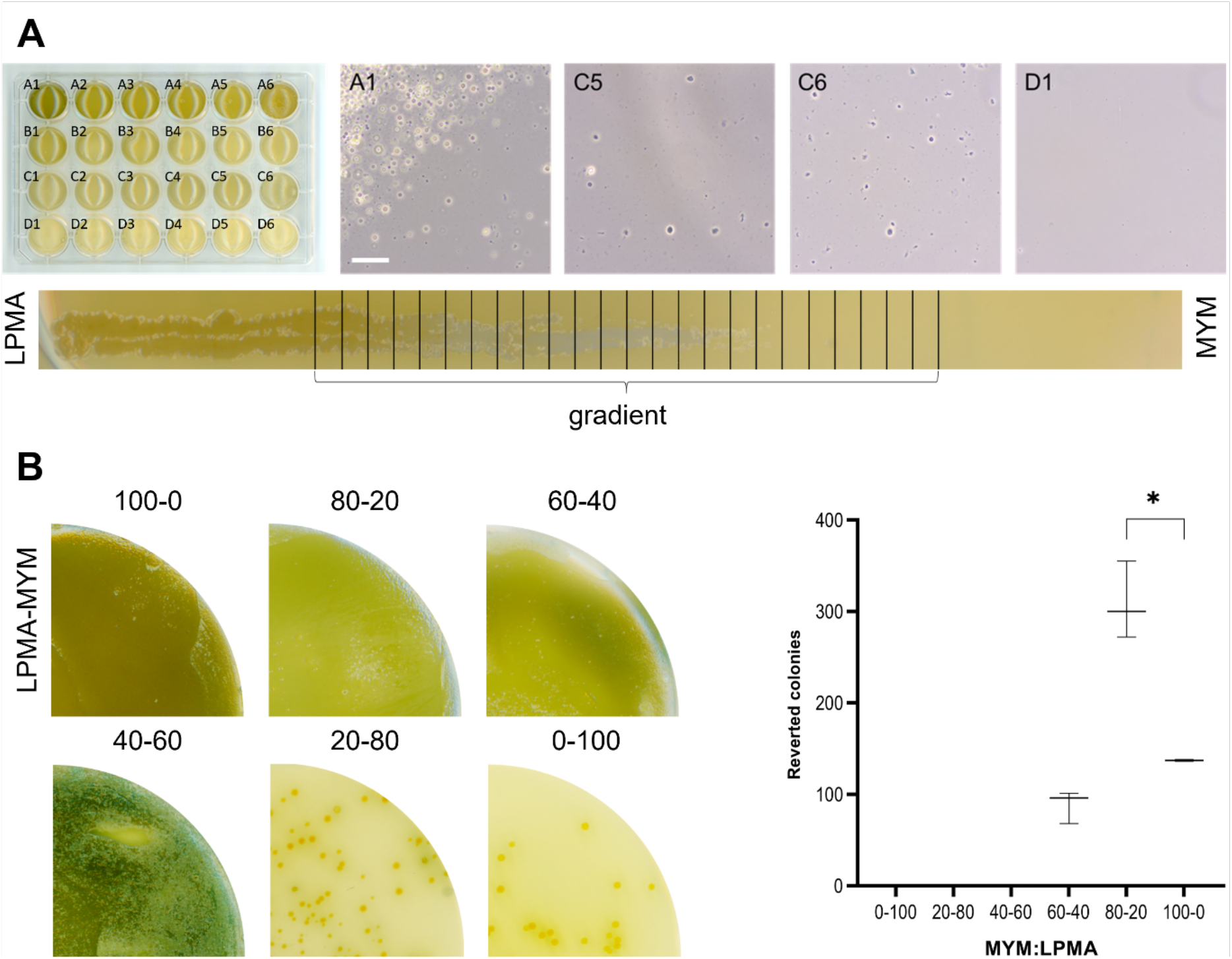
Gradient agar plates allow prediction of switching efficiency. (**A**) Comparison of switching efficiency of Δ*dcw* between 24-well plates containing defined mixtures of MYM/LPMA and gradient agar plates. In both cases, cells were able to grow up to an MYM:LPMA concentration of 75%:25%. The numbers of the microscopy pictures correlate with the wells on the 24-well plate. The scale bar represents 40 µm. (B) Quantification of switching efficiency of *alpha* in different ratios of MYM/LPMA medium. In comparison to pure MYM medium, the switching frequency of *alpha* increases 2-fold in 80:20 MYM:LPMA medium. At a medium composition of 60% LPMA, all colonies consist of wall-deficient cells, while on 40% LPMA, both morphotypes (walled and wall-deficient) are found.

**Figure 3.**
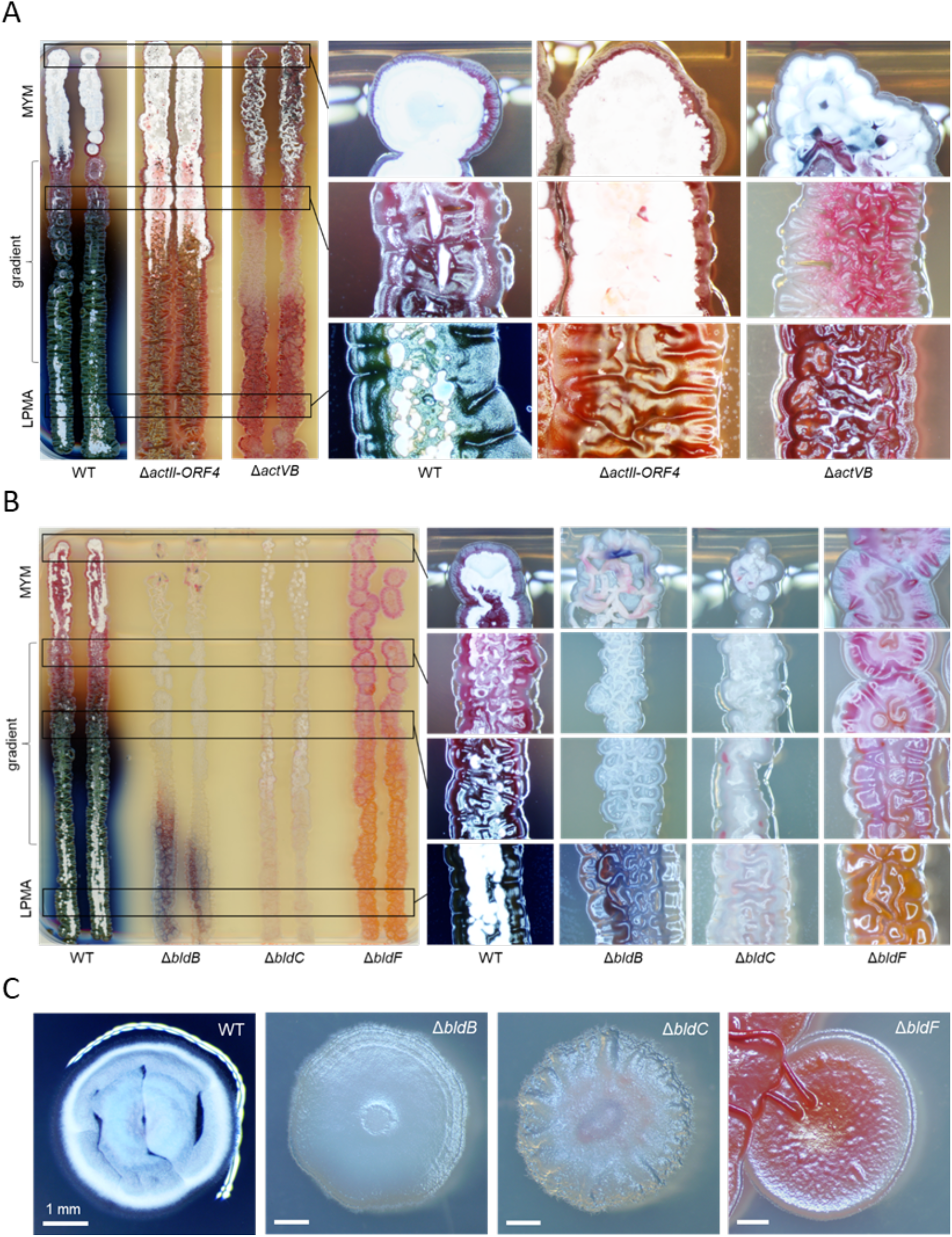
*Streptomyces coelicolor* A3(2) M145 displays both morphological and chemical differentiation on the gradient agar plate. (**A**) Morphological and chemical differentiation of *S. coelicolor* M145 and the actinorhodin-deficient mutants Δ*actVB* and Δ*actII-ORF4*. Please note that an increase in osmolytes leads to an increase in actinorhodin production in the wild-type strain. (**B**) Δ*bldB* and Δ*bldF* were unable to form aerial hyphae, while Δ*bldC* was able to grow aerial hyphae. (**C**) Bld mutants grown on standard R5 medium displayed no antibiotic production nor aerial hyphal growth.

The gradient agar plate experiments also revealed that only a small fraction of *alpha* cells was able to revert to mycelial growth on 0% LPMA medium but suggested that a larger fraction of cells could revert when mixtures of LPMA and MYM were used (Fig. 1). Therefore, we grew *alpha* on six different ratios of LPMA and MYM in triplicate. Indeed, by using a mixture of LPMA/MYM (20:80), more than twice the number of colonies observed were able to switch to mycelial growth (Fig. 2B). A significant difference was found between the number of reverted colonies grown on 40-60 and 20-80 LPMA:MYM, and 20-80 and 0-100 LPMA:MYM (unpaired t-test with Welch’s correction, p = 0,0054 and p = 0,0195, respectively).

### 3.3 Patterns of morphological and chemical differentiation are dependent on osmolyte concentration

To assess the usability of gradient plates for studying morphological differentiation more broadly, we grew a series of wild-type and mutant *Streptomyces* strains on gradient agar plates. The model streptomycete *Streptomyces coelicolor* produced aerial hyphae and spores on MYM, but readily lost the ability to develop when LPMA concentrations increased to above 30% (Fig. 3). Interestingly, this developmental arrest coincided with the formation of a green-blue metallic pigment. Furthermore, the production of the blue-pigmented antibiotic actinorhodin (Act) increased. Indeed, no blue pigment was observed when the actinorhodin-deficient strains Δ*actVB* (lacking the dimerization enzyme for actinorhodin biosynthesis) and Δ*actII-OR4* (lacking the cluster-specific activator) were used (Fig. 3A). Instead, the Δ*actII-OR4* mutant produced red-pigmented prodiginines (Red) when subjected to osmotic stress, as seen by the distinctive orange coloration. We also grew three *bld* mutant strains, which were previously found to be deficient in aerial hyphae formation on R5 medium (Eccleston et al., 2002; Hunt et al., 2005; Puglia et al, 1984) (Fig. 3C). As expected, Δ*bldB* and Δ*bldF* were unable to form aerial hyphae on the gradient plates, while Δ*bldC* produced some sparse aerial hyphae at 25-100% MYM (Fig. 3B). Notably, Δ*bldB* had a purple pigmentation at 95-100% LPMA, which may represent a combination of Act and Red. In addition, Δ*bldF* produced a bright orange pigment at a percentage of 60-100% LPMA which could be Act (Bystrykh et al., 1996). Altogether, these results show that gradient agar plates can also be used for studying morphological differentiation more broadly amongst Actinobacteria.

## 4. CONCLUSION

As gradient agar plates can be used with a plethora of substances, it is surprising that only a few applications have been explored to date. Examples are pH gradients (Sacks, 1956) or gradients of an antibiotic (Quiblier et al., 2011; Thomas et al., 1996). We have used gradient agar plates as a fast and reproducible method to deduce switching efficiencies. Using agar plates with a gradient of osmoprotectants, strains can gradually revert to a walled state, thereby dramatically decreasing lysis. Furthermore, our method allows precise determination of the osmolyte concentration where reversion takes place, allowing careful and reproducible comparison between mutants. Besides, we show that gradient agar plates can also be applied for another purpose, namely to study chemical differentiation in streptomycetes as a response to osmotic stress. Furthermore, gradient plates could also be used to determine responses to other ecological triggers, such as iron or copper availability, or the presence of lysozyme (Benachour et al, 2012). As bacteria have developed numerous chemical and morphological responses to osmotic stress, agar plates with a gradient of osmolytes have a huge potential to better understand these important stress responses and unravel the underlying mechanisms.

*K. viridifaciens* Δ*dcw* was found to be unable to revert to the walled state, caused by the absence of a large part of the *dcw* cluster (Zhang et al., 2021). Interestingly, our gradient plates reveal colonies at an LPMA percentage of 20% LPMA, indicating that these L-forms could tolerate a much lower concentration of osmolytes than previously thought. Besides, as *alpha* and M1 already revert to a walled state at this medium composition, suggesting that a walled state is the preferred mode of growth even though wall-deficient growth would still be possible. Future work could help us better understand this physiological decision-making.

## Acknowledgments

We would like to thank Véronique Ongenae (Institute of Biology, Leiden University, the Netherlands) for providing feedback on the manuscript. We are grateful to Helga van der Heul (Institute of Biology, Leiden University, the Netherlands) for providing us with the *S. coelicolor* strains. Finally, we would like to thank Le Zhang (Institute of Biology, Leiden University, the Netherlands) for the fruitful discussions.

## Author contributions

Maarten Lubbers: Conceptualization; Data curation; Formal analysis; Investigation; Methodology; Visualization; Writing - original draft. Gilles van Wezel: Methodology; Resources; Supervision; Writing - review & editing. Dennis Claessen: Conceptualization; Funding acquisition; Project administration; Resources; Supervision; Writing - review & editing.

## Supplementary figures

**Supplementary figure 1.**
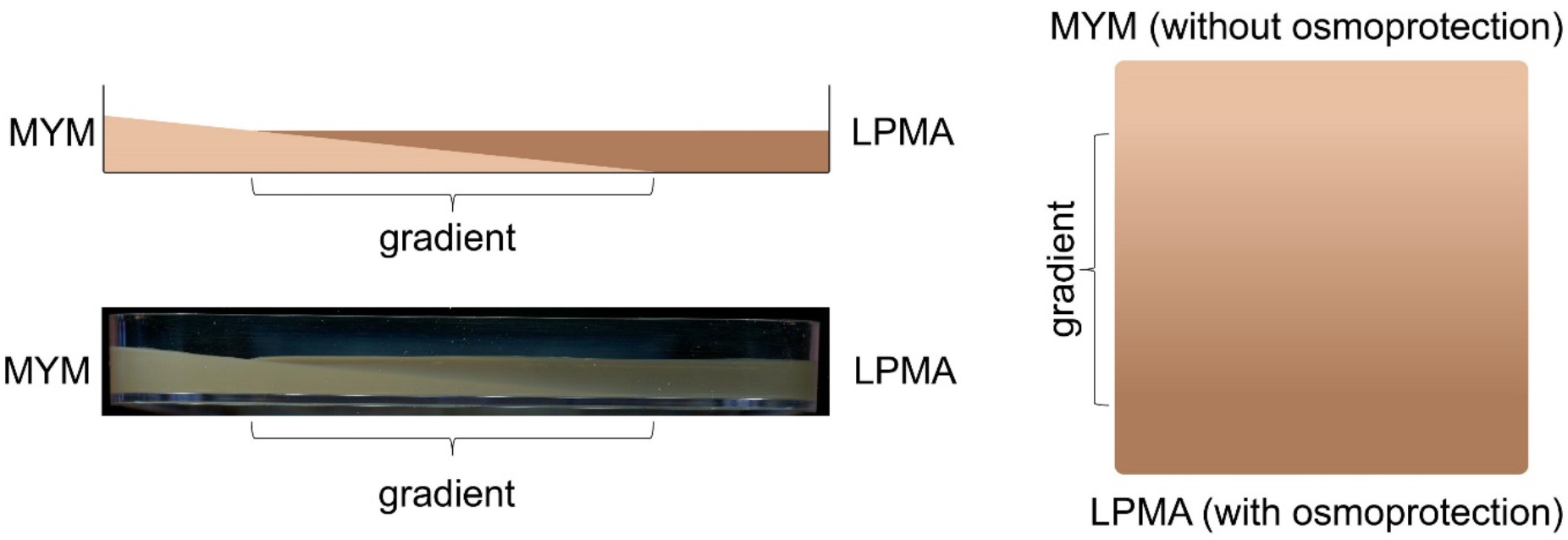
Schematic overview of the gradient agar plates developed for this study, consisting of an inclined layer of MYM with a layer of LPMA on top.

## REFERENCES

1. Benachour, A., Ladjouzi, R., Le Jeune, A., Hébert, L., Thorpe, S., Courtin, P., … & Mesnage, S. (2012). The lysozyme-induced peptidoglycan N-acetylglucosamine deacetylase PgdA (EF1843) is required for Enterococcus faecalis virulence. Journal of bacteriology, 194(22), 6066–6073.

2. Bystrykh, L. V., Fernández-Moreno, M. A., Herrema, J. K., Malpartida, F., Hopwood, D. A., & Dijkhuizen, L. (1996). Production of actinorhodin-related” blue pigments” by Streptomyces coelicolor A3 (2). Journal of bacteriology, 178(8), 2238–2244.

3. de Maayer, P., Anderson, D., Cary, C. & Cowan, D.A. (2014). Some like it cold: understanding the survival strategies of psychrophiles. EMBO Rep 15, 508-517 (2014)

4. Eccleston, M., Ali, R. A., Seyler, R., Westpheling, J., & Nodwell, J. (2002). Structural and genetic analysis of the BldB protein of Streptomyces coelicolor. Journal of bacteriology, 184(15), 4270–4276.

5. Hunt, A. C., Servín-González, L., Kelemen, G. H., & Buttner, M. J. (2005). The bldC developmental locus of Streptomyces coelicolor encodes a member of a family of small DNA-binding proteins related to the DNA-binding domains of the MerR family. Journal of bacteriology, 187(2), 716–728.

6. Kieser, T., Bibb, M. J., Buttner, M. J., Chater, K. F., & Hopwood, D. A. (2000). Practical streptomyces genetics (Vol. 291, p. 397). Norwich: John Innes Foundation.

7. Lebre, P. H., De Maayer, P. & Cowan, D. A. (2017). Xerotolerant bacteria: surviving through a dry spell. Nat Rev Microbiol 15, 285–29.

8. Maccario, L., Sanguino, L., Vogel, T. M. & Larose, C (2015). Snow and ice ecosystems: not so extreme. Res Microbiol 166, 782-795 (2015).

9. Nguyen, J., Lara-Gutiérrez, J., & Stocker, R. (2021). Environmental fluctuations and their effects on microbial communities, populations and individuals. FEMS microbiology reviews, 45(4), fuaa068.

10. Ongenae, V., Mabrouk, A. S., Crooijmans, M., Rozen, D., Briegel, A., & Claessen, D. (2022). Reversible bacteriophage resistance by shedding the bacterial cell wall. Open Biology, 12(6), 210379.

11. Puglia, A.M., and Cappelletti, E. (1984) A bald superfertile U.V.-resistant strain in Streptomyces coelicolor A3(2). Microbiologica 7: 263–266.

12. Quiblier, C., Zinkernagel, A. S., Schuepbach, R. A., Berger-Bächi, B., & Senn, M. M. (2011). Contribution of SecDF to Staphylococcus aureus resistance and expression of virulence factors. BMC microbiology, 11(1), 1–12.

13. Ramijan, K., Ultee, E., Willemse, J., Zhang, Z., Wondergem, J. A., van der Meij, A., Heinrich, D., Briegel, A., Van Wezel, G.P., Claessen, D (2018). Stress-induced formation of cell wall-deficient cells in filamentous actinomycetes. Nature Communications, 9, 5164.

14. Ramijan, K., van Wezel, G. P. & Claessen, D. (2017). Genome sequence of the filamentous actinomycete Kitasatospora viridifaciens. Genome Announcements 5, no. 6, e01560–16

15. Sacks, L. E. (1956). A pH gradient agar plate. Nature, 178(4527), 269–270.

16. Shitut, S., Shen, M. J., Claushuis, B., Derks, R. J., Giera, M., Rozen, D., Claessen, D., Kros, A. (2022). Generating heterokaryotic cells via bacterial cell-cell fusion. Microbiology Spectrum, e01693–22.

17. Stuttard C. 1982. Temperate phages of Streptomyces venezuelae: lysogeny and host specificity shown by phages SV1 and SV2. Microbiology 128:115–121.

18. Szybalski, W., & Bryson, V. (1952). Genetic studies of microbial cross resistance to toxic agents. 1. Cross resistance of Escherichia coli to 15 antibiotics. J. Bacteriol. 64:489–499

19. Thomas, L. V., & Wimpenny, J. W. (1996). Competition between Salmonella and Pseudomonas species growing in and on agar, as affected by pH, sodium chloride concentration and temperature. International journal of food microbiology, 29(2-3), 361–370.

20. Zhang, L., Ramijan, K., Carrión, V. J., van der Aart, L. T., Willemse, J., van Wezel, G. P., & Claessen, D. (2021). An Alternative and Conserved Cell Wall Enzyme That Can Substitute for the Lipid II Synthase MurG. Mbio, 12(2), e03381–20.

