## Supplementary material for "Reproducible switching between a walled and cell wall-deficient lifestyle of actinomycetes using gradient agar plates": Sup. Fig

**Supplementary figures**

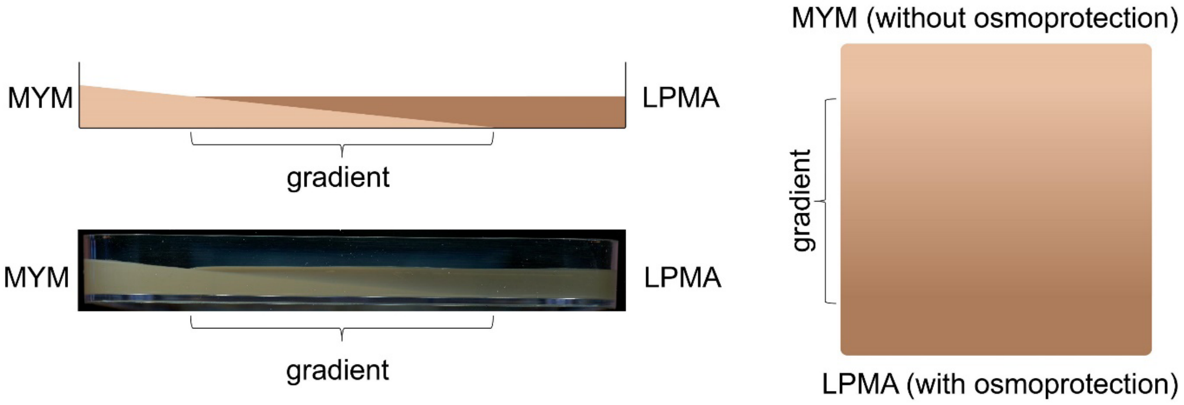


**Supplementary figure 1.** Schematic overview of the gradient agar plates developed for this study, consisting of an inclined layer of MYM with a layer of LPMA on top.
